# Distribution Patterns of rRNA Copy Number Repeats in Prokaryotic Genomes

**DOI:** 10.1101/2025.07.15.664977

**Authors:** Mark Williamson

**Affiliations:** Department of Population Health, School of Medicine & Health Sciences, 1301 N Columbia Rd Stop 9037, University of North Dakota, Grand Forks, ND 58202, United States

**Keywords:** rRNA, copy number, prokaryotes, bacteria, archaea, genomics

## Abstract

Unlike tandem repeats in eukaryotes, ribosomal RNA (rRNA) operons across prokaryotic genomes are widely distributed. Here, I examined the distribution of 16S ribosomal RNA gene copies using all entries from the Ribosomal RNA Operon Copy Number Database with a copy number of 2 or greater, using a metric—normalized range—that allows for comparisons between copy number. Normalized range varied across rRNA copy number, with the greatest distribution of rRNA gene copies in prokaryotic accessions with a copy number of 5. There was a significant phylogenetic signal for both bacteria and archaea (p<0.05). Archaea had higher normalized range than bacteria, even when comparing bacteria with comparable copy number (n=2-5) (p<0.05). Normalized range varied across bacterial phyla (p<0.05), with Cyanobacteria having the largest normalized range and Chlamydiae the smallest. Furthermore, normalized range was predicted by copy number, tRNA number, and pseudo-gene number, after correcting for phylum. When restricted to low bacterial copy numbers (n=2-5), normalized range was predicted by genome size, GC content, tRNA number, coding gene number, and pseudo-gene number. For archaea, normalized range was predicted by genome size, GC content, and tRNA number. Comparisons of bacteria and archaea suggest different mechanisms for rRNA copy distribution and maintenance across genomes.

## 1. Introduction

Ribosomal RNA (rRNA) is a vital component of all living organisms, contributing to the physical structure of the ribosomes to allow them to translate RNA into proteins. Because the 16S component of rRNA in prokaryotes consists of both highly conserved and highly variable regions, it has become a standard tool in the molecular identification of microbes.

The 16S rRNA segment is organized in an operon that typically includes a copy of 5S and 23S rRNA, as well as internally transcribed spacers, all being transcribed together. An important feature of the operon is that it often appears in repeated copies and may be widely spaced across the genome (Vetrovsky and Baldrian, 2013). The phenomenon of having multiple copies is termed copy number (CN). 16S rRNA CN ranges from 1-5 in archaea and 1-21 in bacteria, though the median bacterial CN is 6. These repeats are usually identical or very nearly so, via mechanisms of concerted evolution (Espejo and Plaza, 2018).

Concerted evolution, most evident in rRNA genes (Parks et al., 2019), occurs when genes with multiple copies are conserved within a species but divergent across species (Haig, 2021). In other words, copies do not evolve separately but together (Zhang et al., 2021). Therefore, mutations on one rRNA gene copy are erased, or transferred to all other copies (Espejo and Plaza, 2018). The most likely mechanisms posited are gene conversion—non-reciprocal transfer between homologous sequences—or unequal crossing-over (Zhang et al., 2021, Santoyo and Romero, 2005). The evolution rate of rRNA copies is on par with a single-copy gene due to concerted evolution (Ahn et al., 2020). Concerted evolution is contrasted with birth-and-death evolution, where copies of a gene evolve separately (Zhang et al., 2021). Because rRNA is such a vital component of protein translation, it appears beneficial for organisms to maintain sequence consistency across copies. However, there are sequence divergences across rRNA copies for many species of prokaryotes, which may indicate that some variance can be beneficial (Parks et al., 2019).

Because different species have different CNs, 16S amplicon sequencing of microbial samples cannot accurately resolve cell count abundance without correcting for CN. However, CN is not known for every species, and while there are software and molecular methods to estimate CN (Zhang et al., 2009, Perisin et al., 2016, Angly et al., 2014), copy number correction remains a contentious prospect (Gao and Wu, 2023, Louca et al., 2018, Edgar, 2017, Kembel et al., 2012). Copy number repeats are an important piece of prokaryote life history, as a higher CN is associated with higher potential growth rate or adaptation to environmental changes, while lower levels may confer more growth efficiency (Pereira-Flores et al., 2019, Roller et al., 2016, Valdivia-Anistro et al., 2015, Condon et al., 1992, Stevenson and Schmidt, 2004, Condon et al., 1995). Copy number is important on a community scale and changes across microbial succession and disturbance (Ortiz-Alvarez et al., 2018, Fierer et al., 2010, Nemergut et al., 2016, Guittar et al., 2019, Williamson, 2024).

In eukaryotes, the analogous 18S rRNA operon is typically found in tandem repeats (Hori et al., 2023). In contrast, 16S rRNA copies in prokaryotes can be widely distributed across the circular genome. This raises a fundamental question: how do widely distributed prokaryotic rRNA copies retain their similarity through concerted evolution? If copies are clustered on the genome, concerted evolution mechanisms in operation seem straightforward. But how do widely distributed copies coordinate sequence conservation? Furthermore, how do evolutionary history and genome architecture affect the distribution of rRNA copies across genomes?

Therefore, I was interested in studying the 16S rRNA copy number distribution of prokaryotes, including differences across phyla and genome content. These can be described in the following research questions:

1. How are copy number repeats of the rRNA operon distributed in prokaryotes?
2. What is the phylogenetic signal of CN distribution?
3. Do CN distributions differ across domain (Archaea vs. Bacteria)?
4. Do CN distributions cluster by taxa (for each CN)?
5. Is CN distribution predicted by genome characteristics?

## 2. Materials and Methods

### 2.1 Dataset Creation

#### 2.1.1 rRNA copy number distribution database pipeline

A fasta file composed of 16 rRNA sequences from the most recent version of the Ribosomal RNA Operon Copy Number Database (rrnDB, version 5.8) (Stoddard et al., 2015), was downloaded and unzipped. For each unique accession record (species with multiple rRNA copies had an entry for each copy), the species name, accession number, and CN were extracted and put into a new file using a custom Python script, for a total of 29,344 bacterial and archaeal entries.

For each entry possessing a unique accession number, the name, taxonomy, 16S rRNA copy location along the genome, and total length of the genome was fetched from Entrez using BioPython and compiled into a Python dictionary, with the accession number as the dictionary key. Additional Python scripts were run to obtain GC content, number of tRNAs, number of coding genes, and number of pseudogenes.

For each accession in the dictionary, the start location of each rRNA copy was divided by the total length of the genome and multiplied by 100 to get a **distance by percentage**. Duplicated values were removed, and missing values were obtained by hand by going through the NCBI database.

#### 2.1.2 rRNA copy number distribution statistics

Accessions with two or more rRNA copies were retained for calculating range metrics. Accessions with a single copy number were not considered further. First, the range between the first and last rRNA copy was calculated. Because prokaryote genomes are circular, range could be calculated in two possible scenarios: crossing and non-crossing. In the crossing scenario, the longest piece of the genome was considered the remainder regardless of the start location of the genome sequence. In the non-crossing scenario, the remainder was the combination of the length from the start/end location to the first rRNA copy added to the length from the last rRNA copy to the start/end location. In this analysis, I used the crossing scenarios, as the last rRNA copy might be close to the first copy and thus overestimated range (Figure 1). To calculate range in the crossing scenario, the distance between every copy number distance was tabulated, and then the largest distance was set as the maximum distance. Subtracting the maximum distance from 1.0—the entirety of the genome—gave the range.

**Figure 1:**
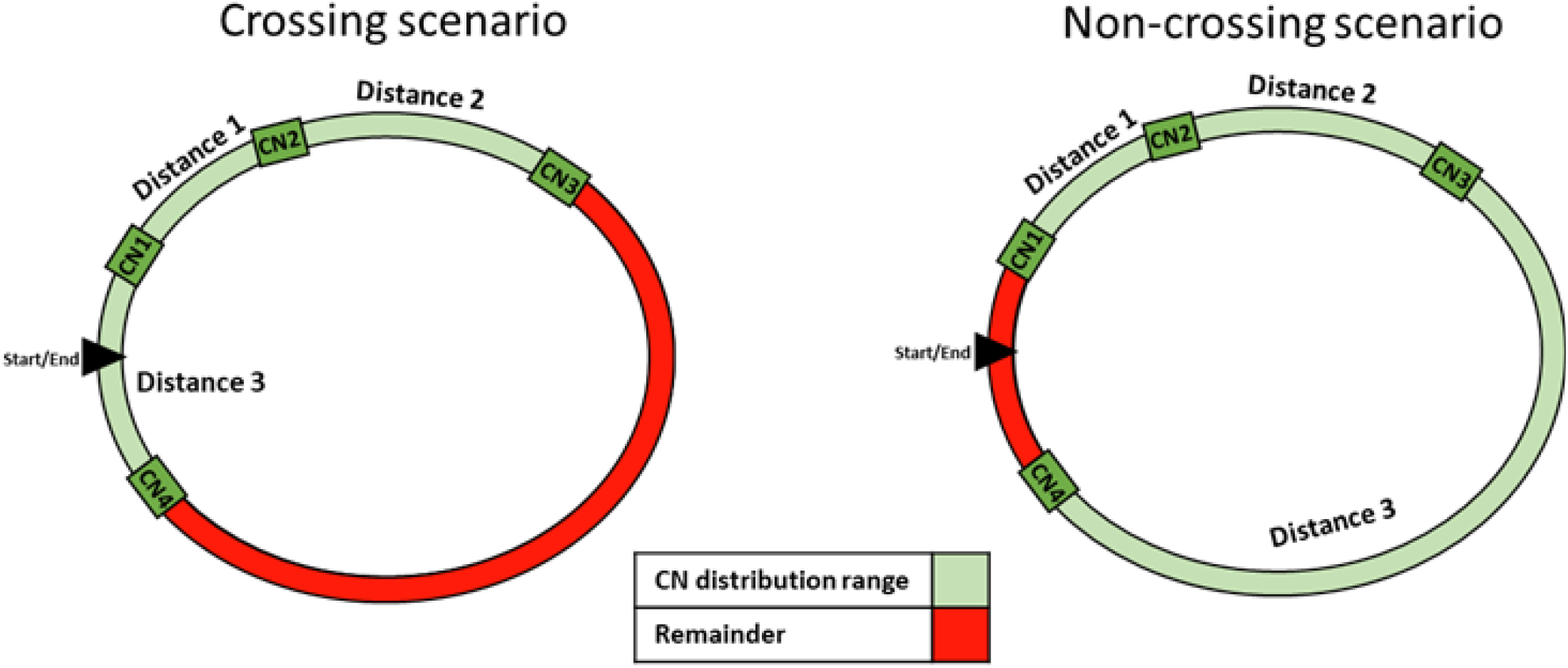
Schematic of rRNA Copy Number Distribution. The circle represents a bacterial genome, with green boxes as rRNA copies. In the crossing scenario, the range is calculated as the total genome minus the section that has the largest remainder, irrespective of the start/end point. In the non-crossing scenario, the range is calculated as the distance between the first (CN1) and last (CN4) rRNA copy.

Accessions with different CNs have different range limits. For example, the range limits for a genome with a CN of two is 0.0 to 0.5, as two rRNA copies can only be a maximum of 50% apart on a circular genome, while the range limits for a genome with a CN of three is 0.0 to 0.66. The lower range limit is always 0.0, while the upper range limit is (CN-1) / CN. To compare accessions across different copy numbers, the range was divided by 1 - (1/CN) to get **normalized range**, with corrected range limits of 0.0 to 0.99 (Box 1). Therefore, I ended up with a metric to compare CN distribution across accessions. However, the gains in interoperability come with a loss in information. Accessions with the same normalized range may have very different internal distributions of additional copies within that range (Figure 2). Furthermore, the location of the first and last rRNA copies can also be widely different. Despite such limitations, the normalized range remains a useful metric of CN distribution that can be compared across prokaryotes.

**Figure 2:**
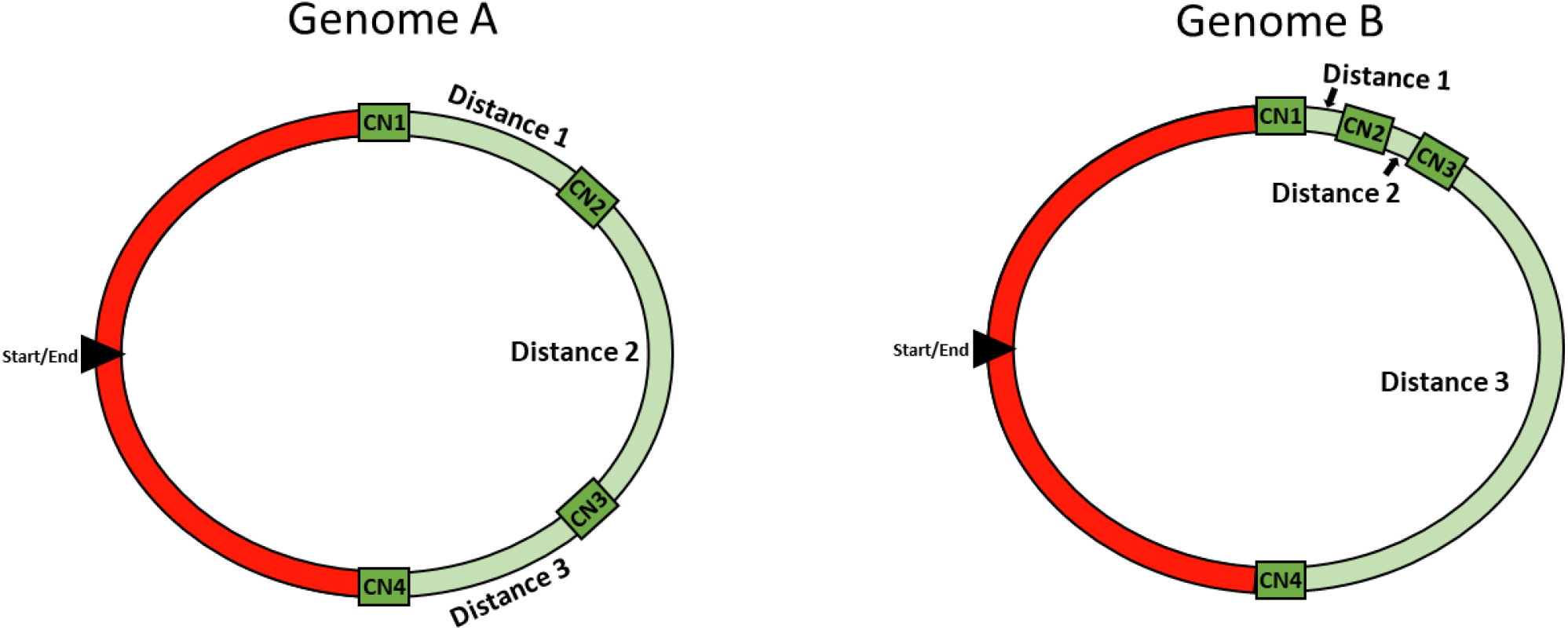
Limitation of normalized range for rRNA Copy Number Distribution. Two genomes with the same normalized range (total distance between first and last rRNA copy is the same) may have very different distributions of the remaining copies. In Genome A, all copies have similar spacing. In Genome B, the first three copies are closely clustered, while the fourth copy is distant.

##### Box 1

**Normalized range calculations**

Example 1. An accession with CN=2, has copies that fall between distances of 0.25 and 0.65.

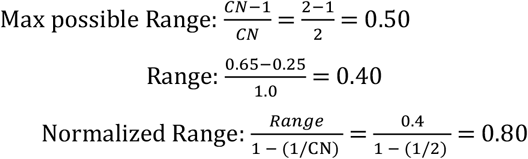

Example 2. An accession with CN=3, has copies that fall between distances of 0.25 and 0.65.

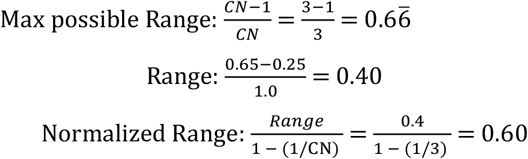

Example 3. An accession with CN=7, has copies that fall between distances of 0.01 and 0.76.

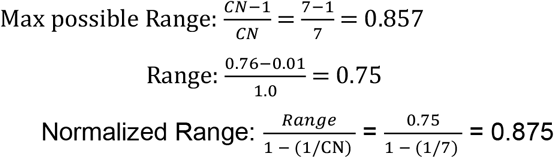

#### 2.1.3 rRNA phylogenetic tree creation

The rrnDB FASTA file of 16s rRNA sequences was split into bacteria and archaea subfiles. Accessions with a single CN were removed. For accessions with multiple rRNA copies, only the first copy was retained for tree creation. The FASTA header was modified to include CN and normalized range.

For the bacterial subfile, the number of sequences was reduced as the large number of accessions made tree construction difficult. First, for species with multiple isolates, only a single entry per species was retained. Second, accessions with CN greater than 10 were removed. Third, ascensions which had no species name (‘sp.’) were also removed. This resulted in 3,979 accessions for bacteria, compared to 193 accessions for archaea. For both FASTA files, bacteria and archaea, trees were aligned using a local version of Clustal Omega 1.2.4 on default settings (Sievers et al., 2011). Trees were created in TreeViewer with default settings and exported as newick files (Bianchini and Sánchez-Baracaldo, 2024).

### 2.2 Data Analysis

To test how copy number repeats of the rRNA operon are distributed in prokaryotes, normalized range of 16S rRNA copy number repeats was plotted using density plots for all prokaryotes, and between domains (bacteria and archaea). Also, density plots were created between bacteria and archaea for accessions constrained to CNs of 2-5, as matching the total CN range for archaea. Normalized range was also compared across CN, with accessions of >=11 CN collapsed into a single group.

To test for phylogenetic signal, bacteria and archaea trees were loaded and checked to ensure that they were binary, ultrametric, and rooted. Trees were then changed to ‘dend’ objects. CN and normalized range were extracted from the header and put into lists. Phylogenetic signal was measured using the function phylosig() from package ‘ape’, using the ‘lambda’ method with CN and normalized range as the signal metrics. Furthermore, because archaea had a sufficiently small set of accessions, a circular phylogenetic tree was created using the package ‘circlize’, and included CN and normalized range information on outer rings.

To test if CN distributions were different across domain, Wilcox tests were run on normalized range between bacteria and archaea. Then, since archaea had a constrained CN range of 2-5, the test was repeated using only bacteria with CN of 2-5. From there, beta regression was run, with normalized range modeled as a function of domain (archaea vs. bacteria) and copy number (2-5).

Next, to test if CN distribution clustered by taxa within domain, bacteria and archaea were examined separately by running a beta regression model with normalized range as a function of phylum. For bacteria, accession frequency across phyla was highly uneven. So, the top ten phyla by frequency count were retained as categories, while the rest of the smaller phyla were grouped into the category ‘Other’ to avoid small frequency cells (Table 1). For archaea, only Euryarchaeota and Thaumarchaeota had enough entries to compare. For significant results, estimated marginal means were generated using the emmeans() function using package ‘emmeans’ and group differences were calculated with a sidak correction.

**Table 1:** Frequency of Phyla in Bacteria and Archaea.

| Phyla | Frequency |
| --- | --- |
| Proteobacteria | 16426 |
| Firmicutes | 6170 |
| Actinobacteria | 2698 |
| Bacteroidetes | 1055 |
| Campylobacterota | 901 |
| Tenericutes | 446 |
| Cyanobacteria/Chloroplast | 206 |
| Spirochaetes | 203 |
| Chlamydiae | 192 |
| Verrucomicrobia | 113 |
| Other | 446 |

Finally, to test if CN distributions were predicted by genome characteristics, beta regressions were run for all prokaryotes, bacteria only, and archaea only, modeling normalized range as a function of genome characteristics. First, simple models with normalized range as a function of copy number as a three-group categorical variable—small (CN=2-4), medium (CN=5-7), and large (CN=8+)—were run. This was followed by simple models with genome size as the predictor variable. For archaeal genome size, there was an outlier with an extremely small genome (accession= NZ_CP031309, name= *Natronorubrum bangense*, strain JCM 10635), that turned out to be a plasmid rather than the chromosome and was removed from the analysis. Then, CN, genome size, GC content, tRNA number, coding gene number, and pseudogene number were rescaled and added as predictor values in a full regression model. The full models tried were both beta regression and Gaussian regression (as normalized range fit both distributions well). From there, a linear mixed model that added phylum as a random effect was run. Lastly, reduced models that removed non-significant variables were run and compared to the full models using AIC scores. The final, best model was determined by AIC score.

## 3. Results

### 3.1 Overall

A total of 29,322 accessions were available from the rrnDB. Of those, 25,644 accessions had a CN of 2 or greater. The majority, 25,450 (99.24%), were bacteria, with 194 (0.76%) remaining for archaea. Copy number ranged from 1-20 in bacteria, and 1-5 in archaea.

The calculated summary of CN distribution, normalized range, was consistent and centered around 0.5, although there were notable subpeaks. Separating by domain, the pattern was recapitulated in bacteria. Archaea peaked close to 0.80 and was heavily left-skewed (Figure 3). Normalized range varied across CN, with the lowest levels in accessions with CN=2, increasing to a maximum when CN=5, and dropping slowly with increasing CN numbers (Figure 4).

**Figure 3:**
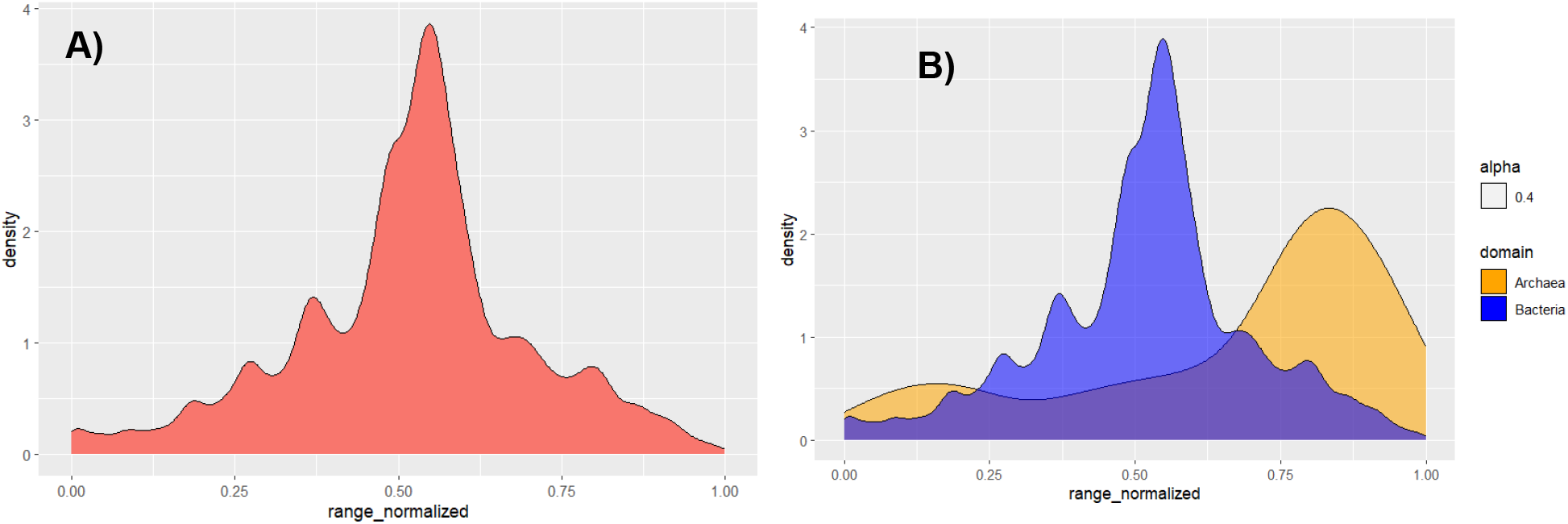
Density Plots of Copy Number Distribution. A) Normalized range across all prokaryote accessions with a copy number of 2 or greater. B) Normalized range across bacteria and archaea separately.

**Figure 4:**
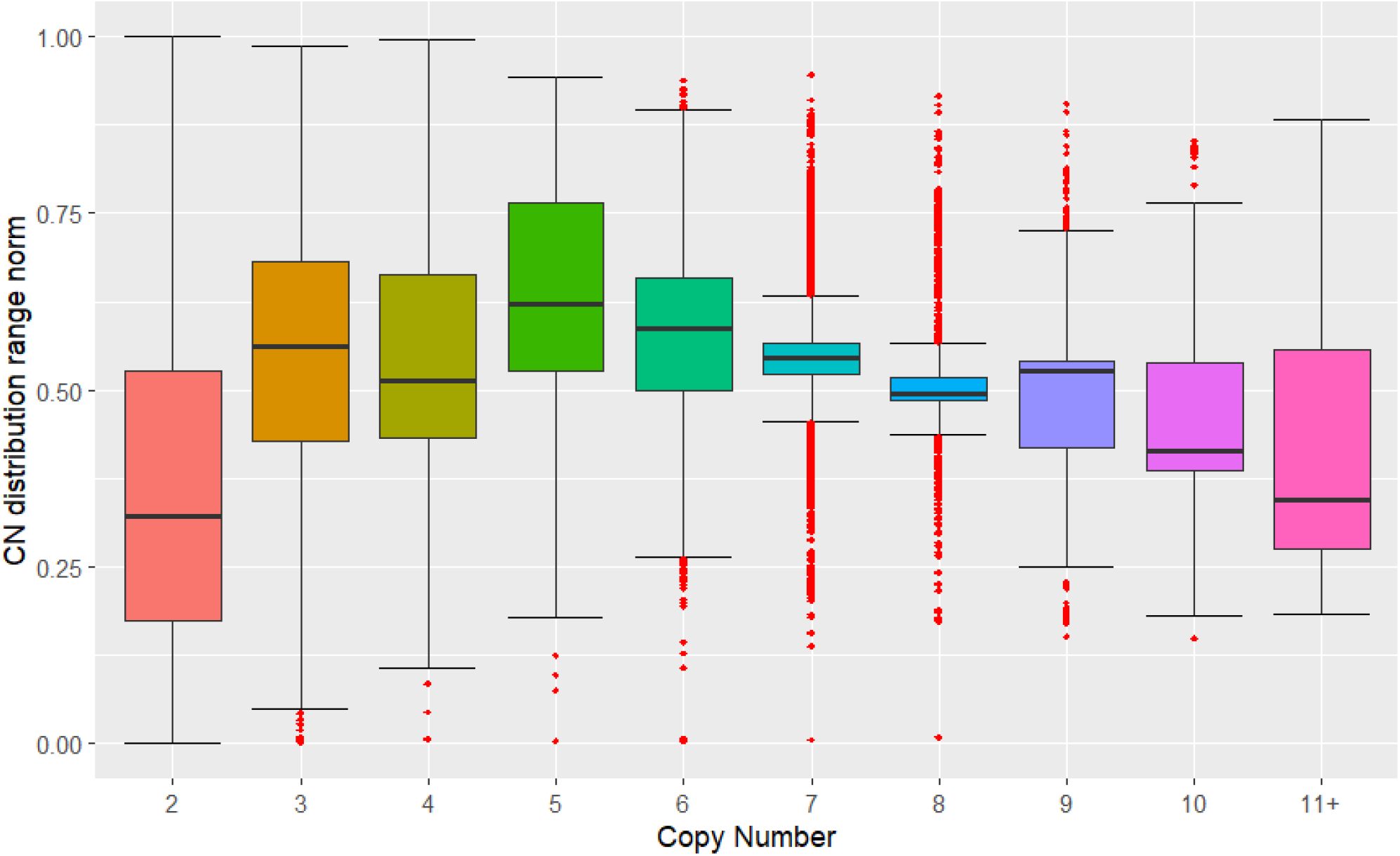
Boxplots of Copy Number Distribution across Copy Number. Normalized range for prokaryotes for copy number category. Highest category (11+) ranges from 11 to 21. Black bar indicates median and red dots indicate outliers.

There was a significant phylogenetic signal for both bacteria (lambda=0.997, p<0.0001) and archaea (lambda=0.995, p<0.0001). Examining a circular plot of a 16S archaeal tree revealed clustering of normalized range in a fine-grain manner, as opposed to the more restrained and homogenous CN variation (Figure 5).

**Figure 5:**
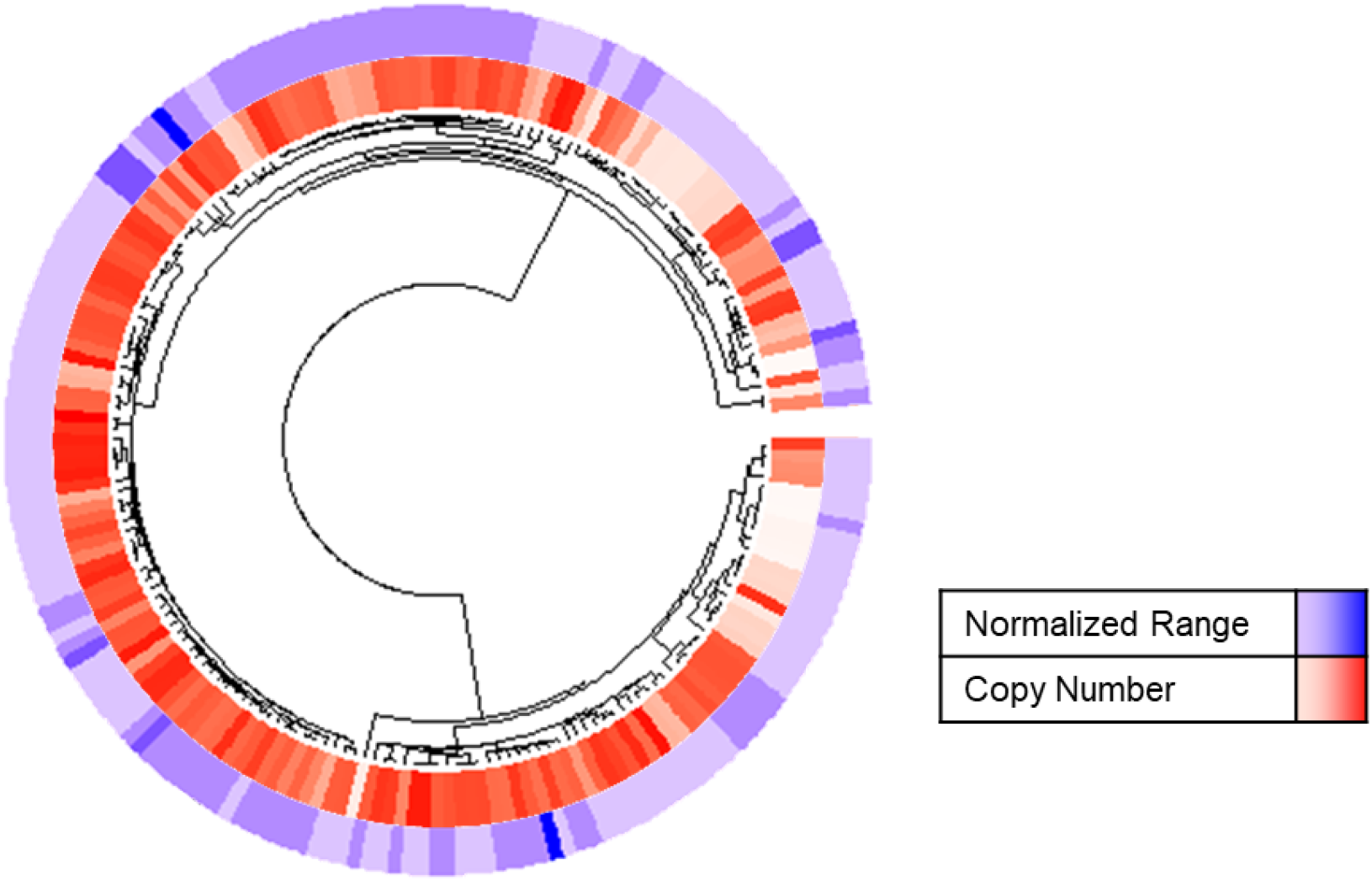
Circular Phylogeny Plots of Archaea. Phylogenetic tree 16S sequences from 194 archaeal accessions, created from a multiple sequence alignment via Clustal Omega and viewed using TreeViewer. The red interior ring indicates copy number of the accessions, ranging from 2 to 5. Blue exterior ring indicates normalized range, ranging from 0.03 to 0.99.

### 3.2 Taxa-Specific

There was a significant difference between normalized range across bacteria and archaea (W=3.5E6, p<0.0001). This remained true when comparing bacterial accessions with CN=2-5 (W=1.6E6, p<0.0001). Normalized range was higher in archaea (mean=0.67, median=0.78) compared to the full bacteria dataset (mean=0.52, median=0.53), and the CN=2-5 reduced dataset (mean=0.51, median=0.52). Normalized range was also significantly different across both domain and CN for the reduced dataset via beta regression (Z=85.6, p<0.0001). Most notably, archaea had consistently higher normalized range across all copy number categories (Figure 6).

**Figure 6:**
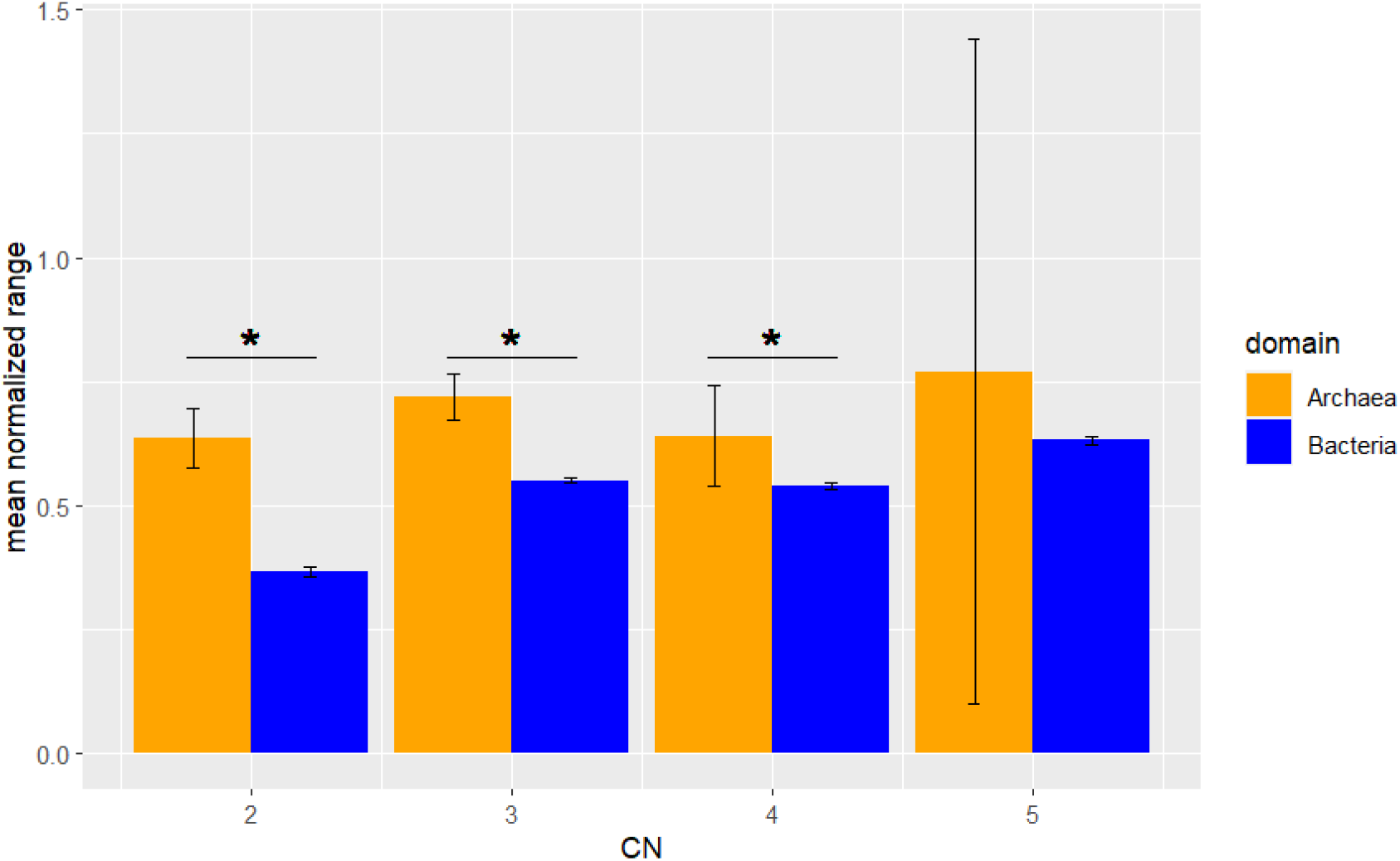
Copy Number Distribution across Domain. Mean normalized range for archaea (orange) and bacteria (blue) that have a rRNA copy number of 2-5. Asterisks indicate significant difference.

Within the bacterial domain, the top ten phyla by accession number were Proteobacteria (Pseudomonadota), Firmicutes (Bacillota), Actinobacteria (Actinomycetota), Bacteroidetes (Bacteroidota), Campylobacterota, Tenericutes (Mycoplasmatota), Cyanobacteria/Chloroplast (Cyanobacteriota, Spirochaetes (Spirochaetota, Chlamydiae (Chlamydiota), and Verrucomicrobia (Verrucomicrobiota). The Other category consisted of 31 different minor phyla. Normalized range was significantly different across bacterial phyla (z-value=121.7, p<0.0005). Based on a post-hoc sidak adjustment, Chlamydiae had the smallest normalized range, noticeably smaller than all other phyla. Cyanobacteria had the highest, along with Bateroidetes, Actinobacteria, and Verrucomicrobia (Figure 7). The difference between Archaeal phyla (Euryarchaeota and Thaumarchaeota) was not significant.

**Figure 7:**
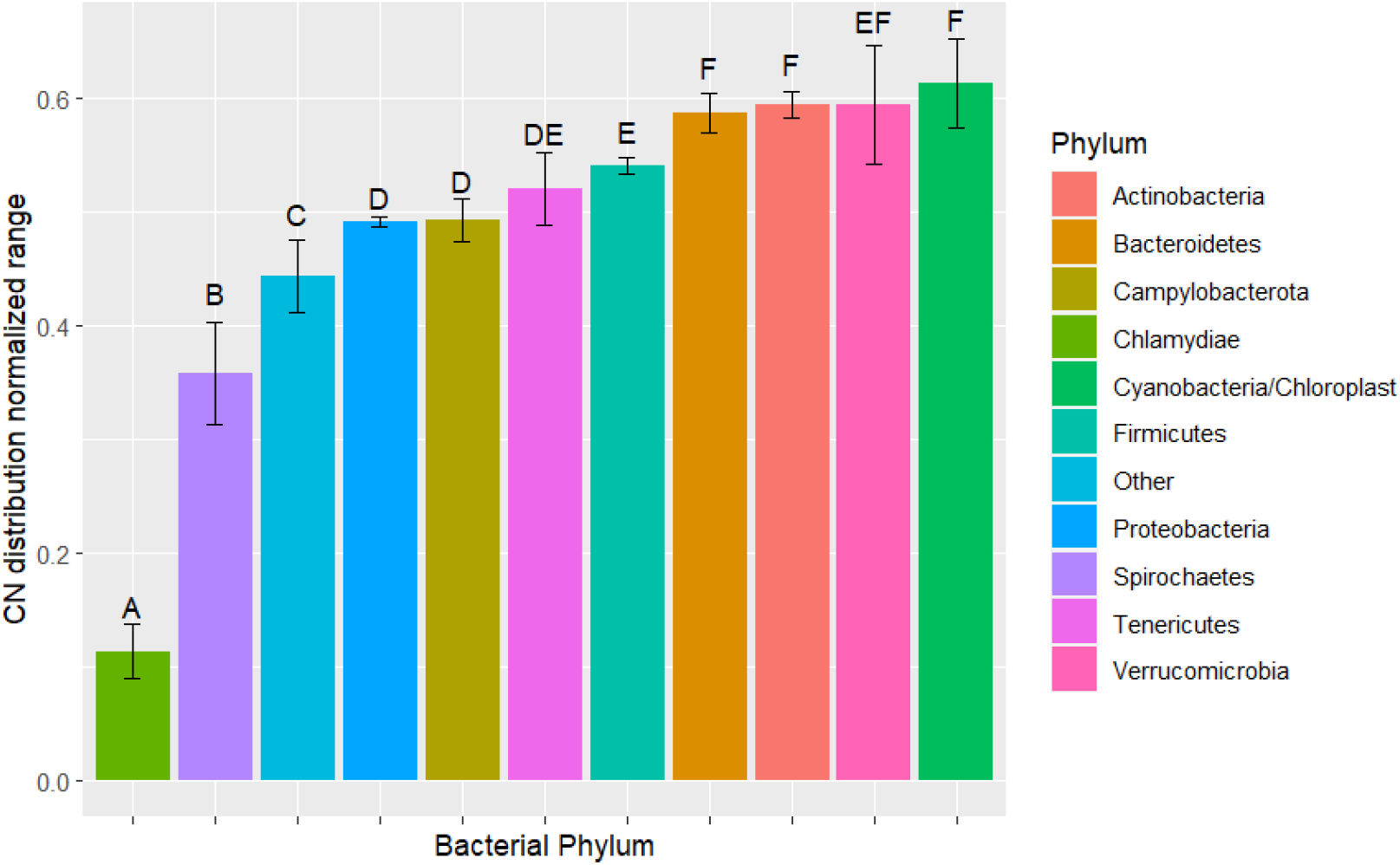
Copy Number Distribution across Bacterial Phyla. Mean normalized range for ten bacterial phyla with highest number of accessions and an ‘other’ category that contains the rest of the smaller phyla. Letters indicate significantly different groups. Phyla with multiple letters were not significantly different from all other groups.

### 3.3 Genome Composition

For models with copy number alone, copy number significantly predicted normalized range in prokaryotes as a whole (Z=12.3, p<0.0001). However, when parsing out bacteria and archaea, the relationship only remained significant in bacteria (Z=10.8, p<0.0001). Testing using a three-group categorical copy number variable (low, medium, high) produced a similar result. The medium category (CN=5-7) had a significantly higher normalized range than low (CN=2-4) and high (CN=8+), with an estimated marginal mean of 0.56 compared to 0.47 and 0.48, respectively (Figure 8a). For models with genome size alone, the pattern was reversed. In bacteria, genome size did not significantly predict normalized range. For archaea, genome size was significant (Z=6.1, p<0.0001). As genome size increased, normalized range increased (Figure 8b).

**Figure 8:**
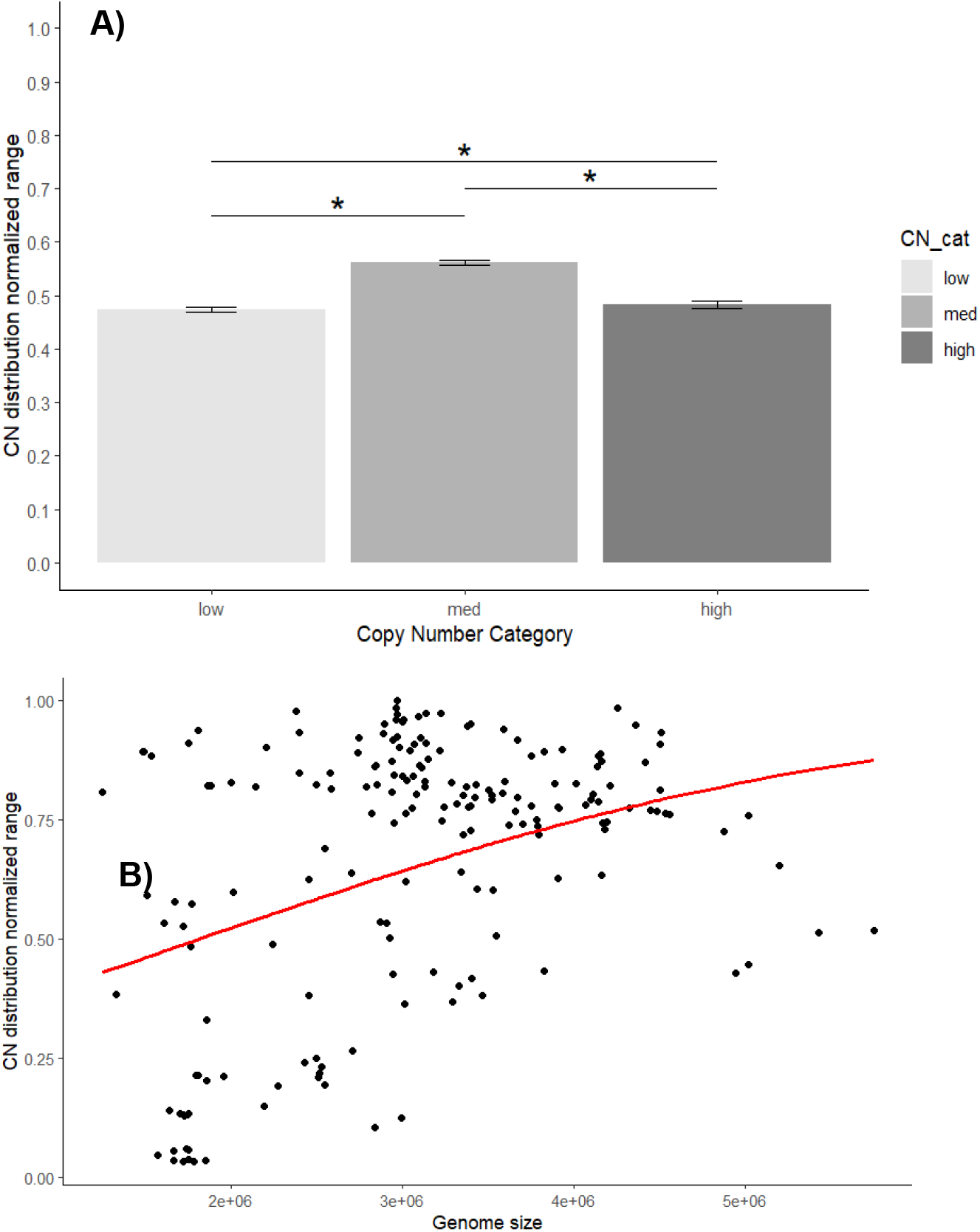
Copy Number Distribution across Copy Number and Genome Size. A) Barplot of mean normalized range across low (2-4), medium (5-7), and high (>=8) copy number in bacteria. B) Scatter plot of normalized range and genome size in archaea, where red line indicates linear model fit.

Given the differences between bacteria and archaea, the two were separated and analyzed individually. Full models with all variables (CN, genome size, GC content, tRNA number, coding gene number, and pseudo-gene number) were run both as a linear model and a linear mixed model with phylum as a random effect.

For bacteria, all variables were significant for the full linear model, but when adding phylum as a random effect, genome size, GC content, and coding gene number were no longer significant. Therefore, a reduced model that included three fixed effect variables—CN, tRNA number, and pseudo-gene number—and phylum as a random effect was also run. Comparing AIC scores between the three models—full linear model, full linear mixed model, reduced linear mixed model—found that the reduced linear mixed model had the best fit (Table 2). CN had a slight, positive correlation with normalized range, while the number of tRNAs and the number of pseudo-genes had slight, negative correlations. As CN increased, normalized range tended to increase. Conversely, when the number of tRNAs or pseudo-genes increased, normalized range tended to decrease. Approximately half the variance in the model was attributed to phylum, justifying including it as a random effect, as well as the previous results that showed a significant difference across bacterial phyla.

**Table 2:** Model Characteristics for Normalized Range and Genome Composition.

| Univariate Models |  |  |  |  |
| --- | --- | --- | --- | --- |
| Groups | Variables | Test statistic | P-value |  |
| Bacteria | CN (numerical) | 12.27 (Z) | <0.0001 |  |
| Archaea | CN (numerical) | 0.270 (Z) | 0.787 |  |
| Bacteria + Archaea | CN (numerical) | 10.80 (Z) | <0.0001 |  |
| Bacteria + Archaea | CN (categorical) | 122.7 (Z) | <0.0001 |  |
| Bacteria | genome size | 0.031 (Z) | 0.9753 |  |
| Archaea | genome size | 6.101 (Z) | <0.0001 |  |
| Bacteria + Archaea | genome size | -0.154 (Z) | 0.878 |  |
| Multivariate Models |  |  |  |  |
| Model | Variables | Test statistic | P-value | AIC Score |
| Full Bacteria generalized linear model | CN | 17.15 | <0.0001 | -15716.18 |
|  | Genome size | 6.63 | <0.0001 |  |
|  | GC content | 2.56 | 0.0105 |  |
|  | tRNAs | -16.79 | <0.0001 |  |
|  | C_genes | -2.61 | 0.0090 |  |
|  | P_genes | -13.879 | <0.0001 |  |
| Full Bacteria generalized linear mixed model | CN | 13.26 | N/A | -17216.39 |
|  | Genome size | 2.17 |  |  |
|  | GC content | -1.41 |  |  |
|  | tRNAs | -12.83 |  |  |
|  | C_genes | -0.66 |  |  |
|  | P_genes | -11.02 |  |  |
|  | Phylum (random) | 0.03 (var) |  |  |
|  |  | 0.03 (resid) |  |  |
| Reduced Bacteria generalized linear mixed model | CN | 15.88 | N/A | -17246.98* |
|  | tRNAs | -13.82 |  |  |
|  | P_genes | -10.39 |  |  |
|  | Phylum (random) | 0.03 (var) |  |  |
|  |  | 0.03 (resid) |  |  |
| Full Archaea generalized linear model | CN | 0.66 | 0.5089 | -11.07 |
|  | Genome size | 2.26 | 0.0247 |  |
|  | GC content | 4.34 | <0.0001 |  |
|  | tRNAs | 2.44 | 0.0155 |  |
|  | C_genes | -1.64 | 0.1032 |  |
|  | P_genes | 0.91 | 0.3620 |  |
| Full Archaea generalized linear mixed model | CN | 0.57 | N/A | 24.34 |
|  | Genome size | 2.27 |  |  |
|  | GC content | 4.04 |  |  |
|  | tRNAs | 2.39 |  |  |
|  | C_genes | -1.43 |  |  |
|  | P_genes | 0.74 |  |  |
|  | Phylum (random) | 0.02 (var) |  |  |
|  |  | 0.05 (resid) |  |  |
| Reduced Archaea generalized linear model | Genome size | 2.06 | 0.0045 | -13.96* |
|  | GC content | 4.55 | <0.0001 |  |
|  | tRNAs | 2.42 | 0.0166 |  |
| Full Bacteria (CN constrained between 2-5) generalized linear mixed model | CN | 37.11 |  | -3899.123* |
|  | Genome size | -12.54 |  |  |
|  | GC content | 7.52 |  |  |
|  | tRNAs | -7.58 |  |  |
|  | C_genes | 10.60 |  |  |
|  | P_genes | -5.26 |  |  |
|  | Phylum (random) | 0.03 (var) |  |  |
|  |  | 0.04 (resid) |  |  |
| Reduced Bacteria (CN constrained between 2-5) generalized linear mixed model | CN | 34.25 |  | -3718.29 |
|  | tRNAs | -1.70 |  |  |
|  | Pgenes | -4.54 |  |  |
|  | Phylum (random) | 0.02 (var) |  |  |
|  |  | 0.04 (resid) |  |  |

For archaea, genome size, GC content, and tRNA number were significant in the full linear model, a result recapitulated in the linear mixed effects model. That result, and because adding phyla as a random effect captured less than a third of the remaining variance, a final model that included the significant variables and no random effects was run, resulting in the best AIC fit (Table 2). Genome size, GC content, and tRNA number all had positive correlations with normalized range. As genome size, GC content, and tRNA numbers increased, normalized range tended to increase as well.

Given the stark differences in the final models between bacteria and archaea, I also ran linear models on bacterial accessions restricted to those with CN of 2-5 to test if major differences were driven by species with higher copy number. For the full linear mixed effects model, all variables appeared significant. Comparing a reduced linear mixed effects model with the same variables as the full bacteria dataset (CN, tRNA number, and pseudo-gene number) was a worse fit via AIC than the full model. This meant for normalized range in bacteria with 2-5 CN, significant predictors were CN, genome size, GC content, tRNA number, coding gene number, and pseudogene number. For prokaryotes broadly, there were no shared genome characteristics between all bacteria and archaea. Bacteria limited to 2-5 CN overlapped, having significant characteristics of both.

## 4. Discussion

Copy number distribution, as measured by normalized range, varied across prokaryotes and exhibited strong phylogenetic signal. It increased across copy number, with the highest levels at in accession with a copy number of 5, decreasing at higher copy numbers. Archaea had consistently higher normalized ranges across similar CN categories compared to bacteria. Normalized range also varied across bacterial phyla. Normalized range was associated with genome features. It was positively associated with CN and negatively associated with tRNA and pseudogene number in bacteria, while it was positively associated with genome size, GC content, and tRNA number in archaea.

Normalized range is a useful measure of the distribution of 16S rRNA copies in a genome that can be compared across genomes with different numbers of copies. A value of 1.0 means maximum separation, equally spaced copies across the circular genome. A value near 0.0 means all the copies are tightly clustered. Bacteria have on average a normalized range of slightly above 0.5. A genome with two 16S copies would have a 25% separation. However, for higher copy numbers, the pattern is more complicated, as the same normalized range could result in many different patterns. For example, a genome with 3 copies may have two copies closely clumped with a third copy highly spaced, while in another genome all three copies are spaced evenly with the same range. Within the distribution of bacterial normalized range, there are several sub-peaks, more noticeably on the lower end. This may indicate that there are more stable configurations across normalized range. Separating by CN for bacterial accessions with 2-5 copies, showed a complex pattern of peaks (Supplemental Figure 1a), without any clear interpretation.

The density plot of normalized range for archaeal accessions revealed a higher peak, above 0.80, indicating that copies tend to be maximally dispersed across archaeal genomes. There is a small, rounded peak around 0.15, suggesting that for a few species, clustered copies are present. Looking at the density by copy number category shows a complex pattern, although accessions with a CN of 2 are more evident at lower levels (Supplemental Figure 1b).

Normalized range differed across domain. Archaea had consistently higher values, meaning that the 16s rRNA copies were more spaced out in the genome. This may suggest a diverging mechanism for copy number repeat events and maintenance of copies. However, I was unable to find any studies that specifically looked at the differences between bacterial and archaeal rRNA CN evolution. Bacteria and archaea are considered together in relevant sources (Liao, 2000, Rogers, 2019, Acinas et al., 2004, Santoyo and Romero, 2005).

Within archaea, it was not surprising that normalized range was not different between phyla, as only two—Thaumarchaeota and Euryarchaeota—had enough accessions with CN>=2 to analyze. Even with additional data in the future, I expect archaea to have constrained CN distribution, compared to the more diverse bacterial phyla. Known archaea have a maximum copy number of 5 compared to 21 in bacteria.

Within bacteria, the phylum Chlamydiae has a very low mean normalized range, which may be due to most species being pathogenic, resulting in lower CN and more compact, streamlined genomes. Indeed, the rrnDB indicates that the phylum has a mean CN of 1.7. Furthermore, the average genome size is about a quarter of the bacterial average (1.12M bases compared to 4M bases). Phylum Cyanobacteria—well known as primary producers and pioneer organisms—has the highest normalized range, which also might be due to its genomic history. Comparative genomics within the phylum shows an expansion of gene families in response to environmental shifts and high horizontal gene transfer, especially in marine species (Chen et al., 2021), which may account for the wide distribution of rRNA CN across the genome. However, other major linages had a similarly high normalized range. Other interesting outcomes were Spirochaetes, Proteobacteria, Verrucomicrobia, and Other. Spirochaetes has the second lowest normalized range, while also being another notable pathogenic-filled linage. This suggests a closer look at CN distribution in pathogenic versus non-pathogenic species. However it should be noted that Spirochaetes has an average genome size more in line with bacteria in general (3.39M bases). Proteobacteria, despite having the most accession entries—over 50%—has an incredibly narrow variation in normalized range, despite the huge variation in CN across the phylum (1-21), which warrants more detailed inspection. Verrucomicrobia, a major lineage in soil and water, has a high normalized range and variation, which suggests that bacteria in complex environments might tend towards more distributed rRNA copies. The Other category— comprising nearly 30 smaller bacterial phyla—had less variation than single phyla like Spirochaetes, Verrucomicrobia, or Cyanobacteria, despite such phylogenetic breadth. A more detailed examination of the minor phyla are warranted in the future.

Bacteria and archaea have very different patterns of high-level genomic characteristics (genome size, GC content, gene number, etc.) that contribute to CN distribution, although restricting bacterial to 16S rRNA copy numbers of 2 to 5 mitigated those differences somewhat. For bacteria, CN and phyla were the major predictors. The number of tRNAs and pseudo genes was also significant for the full sample, though tRNA number was not significant in CN 2-5 accessions. It is unclear why fewer pseudo genes or tRNAs would contribute to a higher CN distribution. A positive association of tRNA number and CN distribution was expected, given that one or more tRNAs are typically included in the rRNA operon (Acinas et al., 2004, Condon et al., 1995). The number of tRNA genes within the operon has also been shown to affect the length of the intergenic spacer region (Osorio et al., 2005). Indeed, a positive association between pseudogene number and CN distribution might even be expected, as distance should weaken the effect of convergent evolution and make the formation of pseudogenes from gene copies such as rRNA more likely. More work needs to be done to examine CN distribution as a function of 16S rRNA copy divergence and the presence of pseudogenes that are specifically of rRNA origin.

CN was not a significant predictor of normalized range in archaea. Higher genome size tended to have higher CN distribution, an expected outcome. The more bases a genome has, the more room that CN repeats have to spread out. This makes the lack of a statistical association between genome size and CN distribution in bacteria even more striking. In archaea, GC content also appeared to be significant, as it has been associated with a variety of features: *genome size*, especially aerobic, facultative, and microaerophilic prokaryotes (Musto et al., 2006, Almpanis et al., 2018); *lifestyle*, with free-living species tending to have higher GC content while nutrient-limited species tending to have lower (Mann and Chen, 2010), and symbionts having unusually large GC content despite a small genome size (McCutcheon et al., 2009); *temperature*, as GC pairs are more stable and appear to be positively correlated with sections of rRNA and tRNA (Galtier and Lobry, 1997), as well as third codons in extremophiles (Kagawa et al., 1984); *mutational pressure*, including major mutator genes as a function of environment and lifestyle (Wu et al., 2012, Muto and Osawa, 1987, Sueoka, 1988); *phylogenetic grouping* (Barcelo-Antemate et al., 2023); *habitat* (Foerstner et al., 2005); and even *protein structure* (Barcelo-Antemate et al., 2023). In contrast to a negative relationship in the full bacteria dataset, archaea had a mild, significant, positive relationship with the number of tRNAs, again more to expectation.

Interestingly, in CN-constrained bacteria (CN=2-5), while genome size was a significant predictor, it was a negative predictor. The average genome size in archaea is 3.04M bases (SD=0.95) while bacteria averages 4.00M bases (SD=1.58) and 3.54M bases (SD=1.71) for CN-constrained bacteria. As genome size increased, CN distribution tended to decrease, further evidence of fundamentally different mechanisms in bacteria and archaea that do not depend simply on CN or genome size.

## 5. Conclusions

This study had several limitations. Inclusion was limited to prokaryote accessions that had CN information in the rrnDB, which overrepresented highly abundant and culturable isolates. Second, given the large size, comparing CN distribution necessarily collapses specific distribution patterns for higher CN into a single metric, normalized range. Third, some grouping variables, such as phyla, were necessarily collapsed to allow for interpretable analyses. Fourth, only CN distribution was examined, not the relative positions of all copy numbers, which may be important for prokaryotic replication as rRNA genes are located near the locus of replication in fast-growing bacteria [29]. Finally, some of the results are difficult to interpret and will require further examination into some of the specific details, such as phyla level.

In contrast, this study has several strengths. It is, to my understanding, the first exploration of the distribution of rRNA copy number across all prokaryotes with CN information. The study also calculated a useful metric, normalized range, that has the same interpretability across varying copy numbers. Future work is needed to understand CN distribution in terms of CN prediction, sequence divergence, sequence convergence, horizontal gene transfer, microbial ecology, gene regions, and human health (ex. differences between pathogenic and non-pathogenic bacteria). In conclusion, CN distribution depends on the number of rRNA copies, exhibits phylogenetic signal, differs between archaea and bacteria, differs between bacterial phyla, and is predicted by different genome features across bacteria and archaea.

## Acknowledgements

This article is dedicated to the memory of my late mentor, Dr. Brian Darby. He instilled in me a love of both statistics and the fascinating world of microbes hidden in plain sight. Requiescat in pace.

## Data availability

The following datasets and code can be found at the GitHub repository: [CopyNumberDistribution](https://github.com/MWilliamson1247/CopyNumberDistribution)

### Datasets

- operon_distribution_prokaryote_2023_master_1way.xlsx -> raw dataset that includes data on individual rRNA genome locations and distance calculations
- M1.csv -> base dataset that includes calculated distribution variables for each accession
- M1_gens.csv -> includes genome size for accessions found in M1
- M4.csv -> includes clear taxonomy for accessions found in M1
- GC_content_gene_number_finale_2024.csv -> includes GC content and gene numbers for accessions found in M1
- Archaea_2023_CN_1plus_16S_rRNA_NORMRANGE.nwk -> phylogenetic tree of archaeal accessions
- Bacteria_2023_CN_1plus_16S_rRNA_rarified_NORMRANGE.nwk -> phylogenetic tree of a subset of bacterial accessions

### Code

- CN_Distribution_Part_1.html -> R Markdown File of the analysis

