## Supplemental Figure 1 for "Distribution Patterns of rRNA Copy Number Repeats in Prokaryotic Genomes"

**Supplemental Figure 1: Density Plots of Copy Number Distribution by Copy Number.** A) Bacterial accessions constrained to a copy number of 2-5. B) Archaeal accessions.

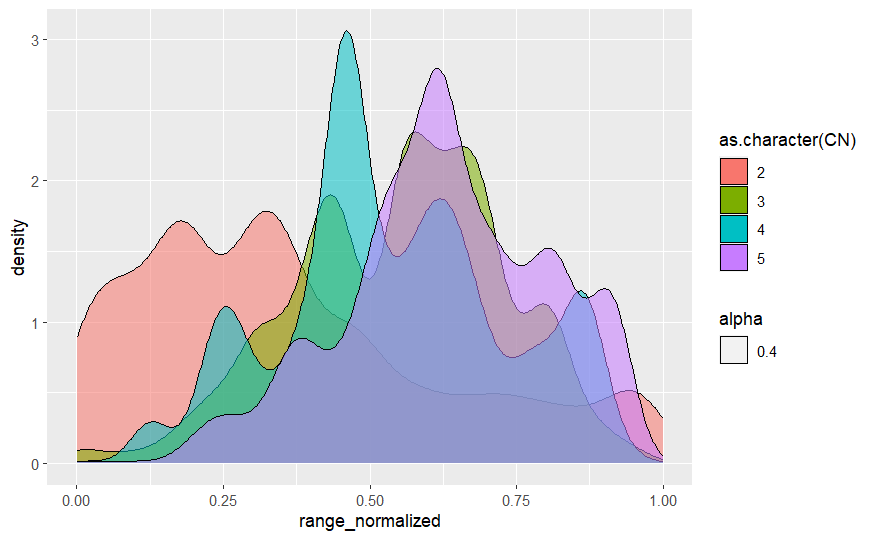

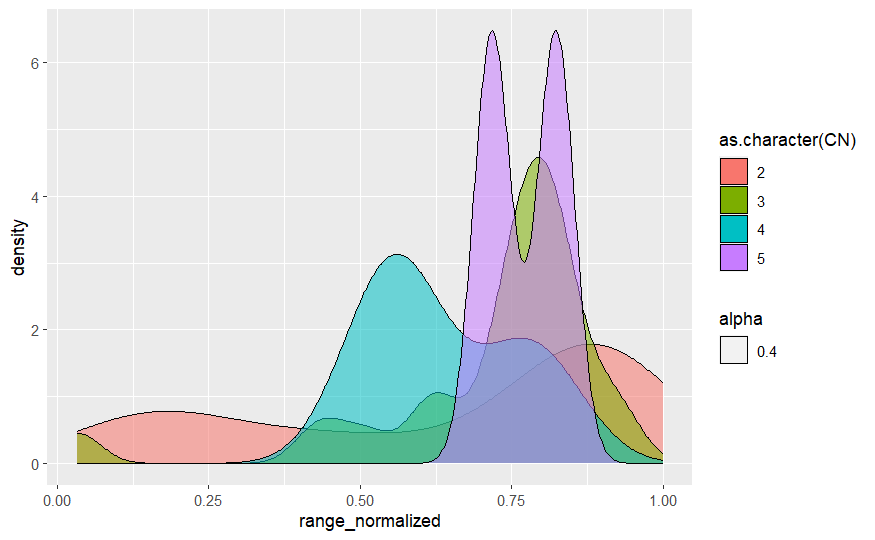

**A)**

**B)**
